# The ancestral shape of the access proton path of mitochondrial ATP synthases revealed by a split subunit-a

**DOI:** 10.1101/2023.02.25.530031

**Authors:** Jonathan E. Wong, Alena Zíková, Ondřej Gahura

## Abstract

The passage of protons across membranes through F_1_F_o_-ATP synthases spins their rotors and drives synthesis of ATP. While the principle of torque generation by proton transfer is known, the mechanisms and routes of proton access and release and their evolution are not fully understood. Here, we show that the entry site and path of protons in the lumenal half-channel of mitochondrial ATP synthases are largely defined by a short N-terminal α-helix of subunit-a. In *Trypanosoma brucei* and other Euglenozoa, the α-helix is part of another polypeptide chain that is a product of subunit-a gene fragmentation. This α-helix and other elements forming the proton pathway are widely conserved across eukaryotes and in Alphaproteobacteria, the closest extant relatives of mitochondria, but not in other bacteria. The α-helix blocks one of two proton routes found in *Escherichia coli*, resulting in the single proton entry site in mitochondrial and alphaproteobacterial ATP synthases. Thus, the shape of the access half-channel predates eukaryotes and originated in the lineage from which mitochondria evolved by endosymbiosis.

## Introduction

F_1_F_o_-ATP synthases are rotary molecular machines that convert the proton-motive force across energy-transducing membranes of bacteria, chloroplasts and mitochondria to the chemical energy of ATP. The flow of protons down their electrochemical gradient through the membrane-intrinsic F_o_ subcomplex spins the enzyme’s asymmetric rotor, whose rotation changes the conformation of three catalytic sites on the membrane-extrinsic F_1_-ATPase subcomplex, resulting in synthesis of ATP. The transmembrane part of the rotor, the c-ring, consists of several copies of subunit-c. Proton translocation is based on sequential protonation of a glutamate residue of each protomer of the c-ring, rotation of the c-ring with the neutralized glutamates exposed to the hydrophobic environment of the phospholipid bilayer, and release of protons on the other side of the membrane. The second element crucial for proton passage is the stator subunit-a (also referred to as ATP6 or subunit-6). Canonical subunits-a contain six α-helices (h_1_-h_6_), five of which are largely embedded in the membrane. While h_1_ is a vertical transmembrane helix, h_3_-h_6_ are horizontal, positioned at an angle ∼20° relative to the plane of the membrane. Helices h_5_-h_6_ form a hairpin bending around the c-ring. The sites of proton binding and release from the c-ring glutamates are separated by a universally conserved arginine of h_5_. The positively charged arginine prevents protons from leaking without c-ring rotation and induces their dissociation at the release site (reviewed in Walker 2017; Kuhlbrandt 2019; Guo and Rubinstein 2022). The proton-binding glutamates on the c-ring are accessible from the surface of the complex by two offset solvent-filled half-channels. In mitochondrial ATP synthases, the access half-channel opens into the lumen of cristae, hence it is called the lumenal half-channel. It is shaped by subunit-a and adjacent membrane elements, including lipids (Klusch, et al. 2017). Based on the high rate of proton translocation, it was proposed that delivery of protons to the c-ring is facilitated by a Grotthuss mechanism through a chain of coordinated water molecules and adjacent amino acid residues (Feniouk, et al. 2004). Recently, the proton-hopping mechanism was supported by observations of cryoEM densities corresponding to water molecules in the inner part of the channel in mammalian (Spikes, et al. 2020) and trypanosomal (Gahura, et al. 2022) mitochondrial ATP synthases. Molecular dynamics simulations are also consistent with the role of coordinated water molecules in proton translocation (Ivontsin, et al. 2022). Direct evidence for the Grotthuss mechanism of proton transfer in both the inlet and outlet channels was provided by a combination of mutagenesis and single-molecule rotation studies with the ATP synthase from *Escherichia coli* (Yanagisawa and Frasch 2021).

Here, we explore the lumenal half-channel forming elements across species and show that the N-terminal region of subunits-a form a short α-helix common to alphaproteobacterial and vast majority of mitochondrial ATP synthases. In Euglenozoa, this helix is a part of a different polypeptide chain encoded by a nuclear gene that is presumably the product of fragmentation of the mitochondrial gene for subunit-a. We document that the helix defines the proton entry site and contributes to the shaping of the inner part of the half-channel in modern eukaryotes and their common ancestors.

## Results and discussion

### The short N-terminal α-helix of subunit-a is a widely conserved structural element of mitochondrial ATP synthases

When examining the proton translocation elements of mitochondrial ATP synthases of two divergent eukaryotes, *Saccharomyces cerevisiae* (Guo, et al. 2017) and *Trypanosoma brucei* (Gahura, et al. 2022), we noticed a remarkable similarity between the N-terminal regions of subunit-a in the yeast and subunit ATPEG3 in the parasite. The N-termini of both proteins form a short single-turn α-helix, hereafter referred to as h_N_, which is located at the layer corresponding to the lumenal surface of the membrane and interacts with the horizontal helix h_5_ of subunit-a in the same manner in both structures. Downstream of h_N_, yeast subunit-a and trypanosomal ATPEG3 follow a similar path; however, after ten residues the yeast chain turns back towards the first transmembrane helix h_1_, thereby diverging from ATPEG3, which continues to the periphery of the complex. The canonical trypanosomal subunit-a, which contains h_1_ and four characteristic horizontal membrane-intrinsic helices h_3_-h_6_, begins nine amino acid residues upstream of h_1_ in close proximity to the site of divergence between yeast subunit-a and ATPEG3 (Fig. 1A). Based on the striking structural and sequence (see below) similarity, we conclude that subunit-a in *T. brucei* has been split and ATPEG3 represents its N-terminal fragment. To acknowledge the origin of ATPEG3, we introduce an alternative name subunit-a_N_.

**Figure 1.**
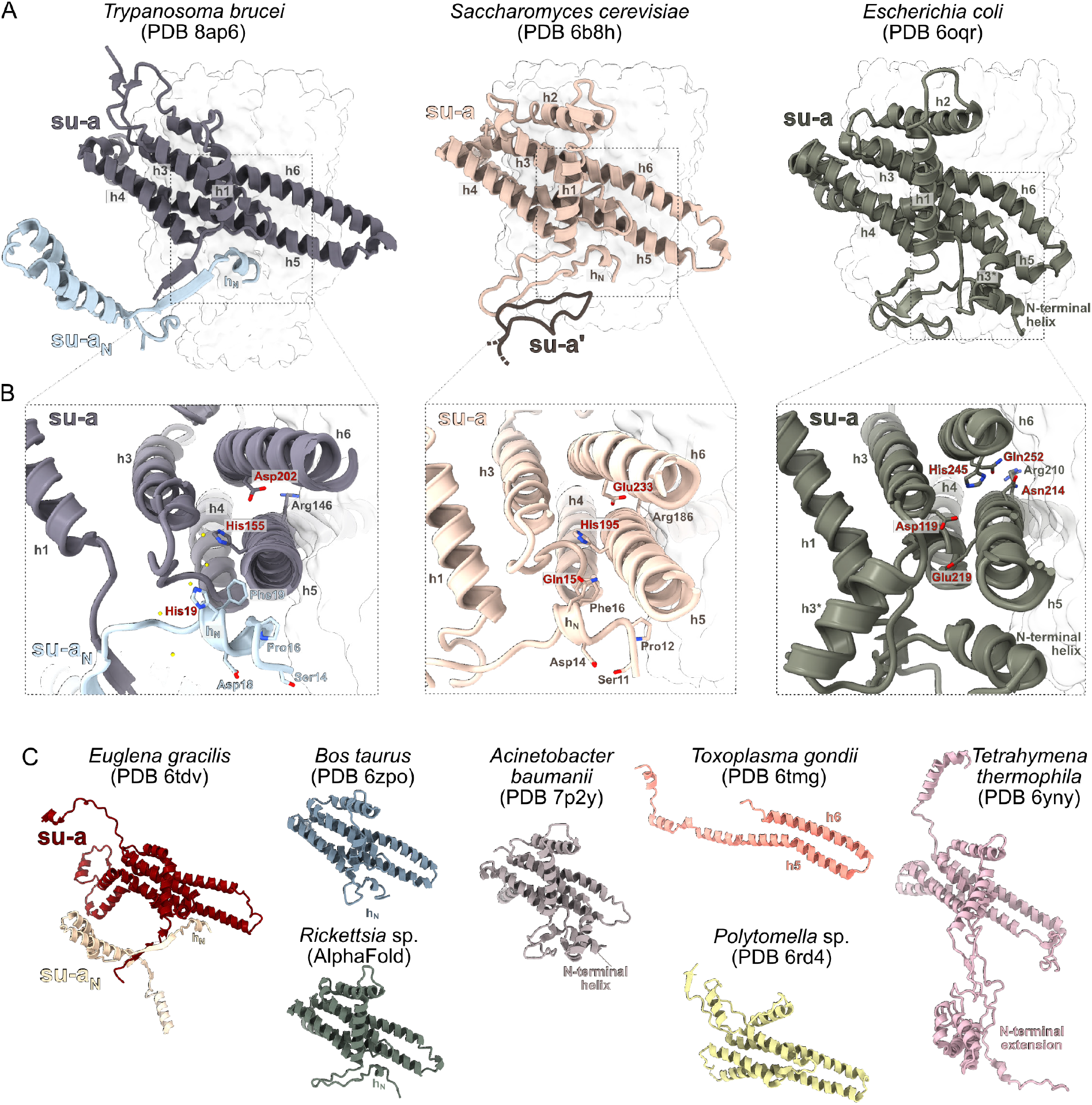
Split subunit-a of mitochondrial ATP synthase in *Trypanosoma brucei* retains ancestral structure and shapes the access proton half-channel. **(A)** Comparison of the structures of subunit-a in *S. cerevisiae* and *E. coli* and subunits a and a_N_ in *T. brucei*. The c-rings are shown as transparent surface. In the structure from *S. cerevisiae*, a fragment of the subunit-a from the other ATP synthase protomer is shown (su-a’). **(B)** The close-up view of the structures in (A). Conserved residues of h_N_ and other selected conserved residues are shown as sticks. The residues labeled in red are directly involved in proton translocation or coordinate water molecules in the Grotthuss chain. Water molecules inferred from cryoEM density are shown as yellow balls in the *T. brucei* structure. **(C)** Structures of subunit-a of mitochondrial and bacterial ATP synthases from selected species. The structure from *Rickettsia* was predicted by AlphaFold.

The split of subunit-a in *T. brucei* occurred in the region corresponding to the loop involved in ATP synthase dimerization in *S. cerevisiae* (Guo, et al. 2017). In *T. brucei*, the loop is structurally replaced by a two-stranded β-sheet, formed by the N-terminus of subunit-a and a β-strand of subunit-a_N_ (Fig. 1A,B). Unlike in yeast, this structure does not contribute to the ATP synthase dimerization interface, which is provided by subunits g and e in this organism (Gahura, et al. 2022). After splitting, subunit-a_N_ was extended at its C-terminus. The extension interacts exclusively with lineage-specific subunits (ATPTB1, ATPTB6, ATPTB11, ATPTB12) or with lineage-specific extensions of subunits shared with other ATP synthases (subunit-i/j and k; Gahura, et al. 2022). It extends to the periphery of the membrane part of the F_o_-moiety, where it co-constitutes a euglenozoan-specific subcomplex, separated from the conserved part by a lipid-filled cavity (Muhleip, et al. 2019; Gahura, et al. 2022). Consistently, fragmented subunit-a is also discernible in the structure of ATP synthase from *Euglena gracilis* in contrast to other mitochondrial and bacterial subunit-a exemplified by several selected species (Fig. 1C; Muhleip, et al. 2019; Gahura, et al. 2021).

The sequence of h_N_ and flanking residues in yeast is highly similar to the N-terminal sequence of subunit-a in all major groups of Alphaproteobacteria (Fig. 2, Supplementary table S1 and Supplementary data 1&2), the closest extant relatives to mitochondria (Munoz-Gomez, et al. 2022). We predicted the structure of subunit-a from *Rickettsia rickettsii* using AlphaFold (Jumper, et al. 2021). The model resembles structures of bacterial and mitochondrial homologs resolved experimentally and includes h_N_ located at exactly the same position as the h_N_ in *T. brucei* and *S. cerevisiae* (Fig. 1C). Similarity of this region in ATP synthases from mitochondria of two deeply branching eukaryotic lineages and the group of bacteria from which mitochondria evolved strongly indicates that the observed architecture of the N-terminal segment of subunit-a is ancestral.

**Figure 2.**
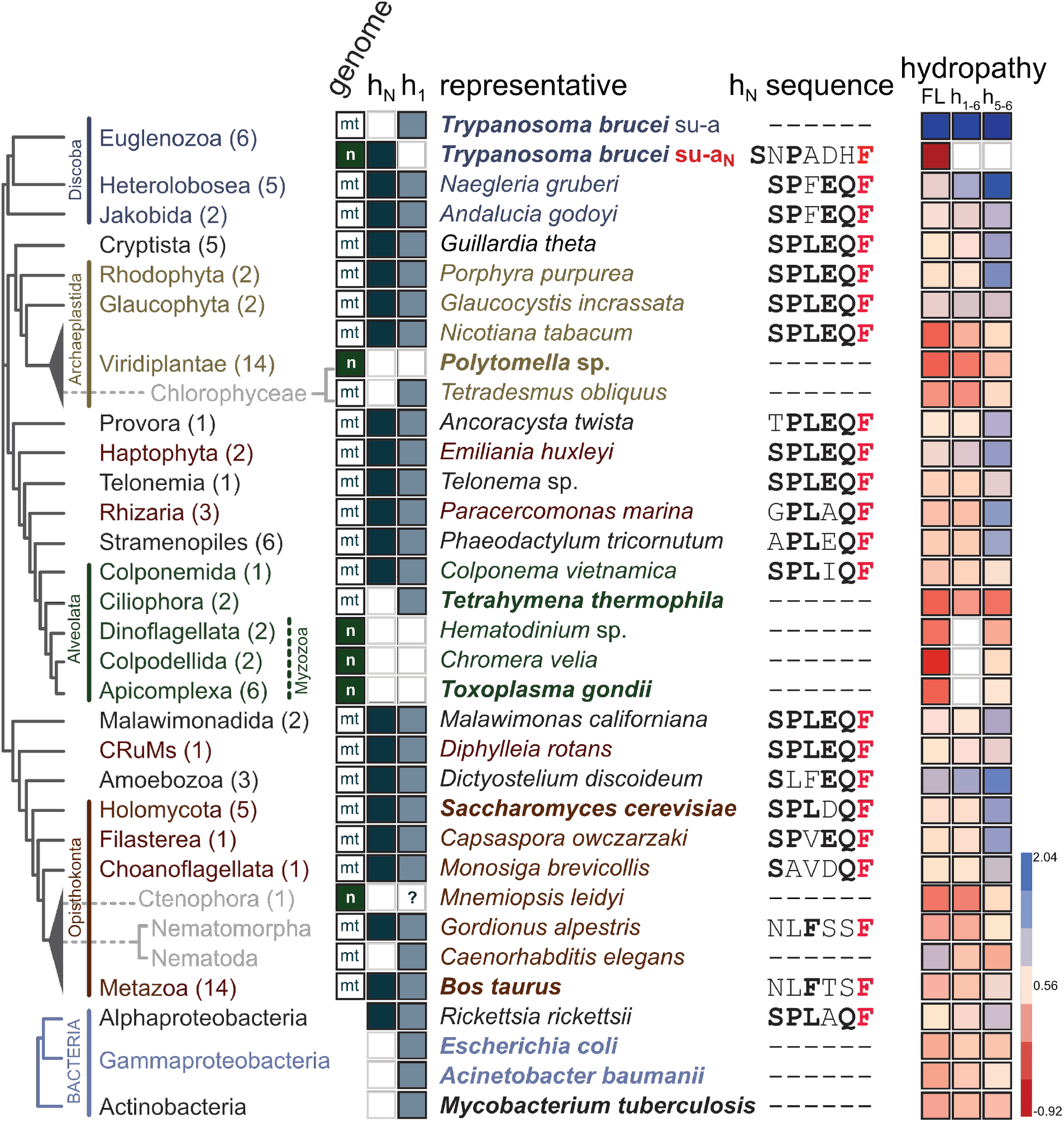
Helix h_N_ is conserved in most mitochondrial ATP synthases and Alphaproteobacteria. For individual lineages mapped on the phylogenetic tree based on (Burki, et al. 2020; Tikhonenkov, et al. 2022) following features are shown: number of species included in the analyses (in parentheses); genome, in which subunit-a/a_N_ is encoded (“mt” – mitochondrial, “n” – nuclear); presence or absence of helices h_N_ and h_1_ of subunit-a/a_N_; selected representative species used in the analyses presented in the following columns (for species shown in bold a cryoEM structure of ATP synthase has been reported); sequence of h_N_ and flanking residues (universally conserved phenylalanine is shown in red, other conserved residues of the SPLEQF motif, or the FXXF motif in Metazoa, are shown in bold); average hydrophobicity of the entire subunit-a (FL), region h_1_-h_6_, and region h_5_-h_6_ (blue – more hydrophobic, red – less hydrophobic).

We compared subunit-a sequences from 94 species covering all major eukaryotic lineages, and structures from selected species determined by cryoEM or predicted by AlphaFold. The sequence comparison documented that the split of subunit-a observed in *T. brucei* and *E. gracilis* is restricted to Euglenozoa, because subunits-a from all representatives of the sister lineage Heterolobosea contain a conserved sequence of h_N_. Consistently, the subunit-a_N_ was found in euglenids, diplonemids and kinetoplastids, the three lineages of Euglenozoa (Sinha and Wideman 2023; Supplementary table S1, Supplementary data 3). Further, the analysis revealed high sequence similarity in the N-terminal region, with most sequences showing a degenerate sequence motif SPLEQF (Fig. 2, Supplementary table S1), supporting the notion that presence of h_N_ represents an ancestral state. The only organisms without the SPLEQF motif were most alveolates, chlorophycean algae *Polytomella* and *Chlamydomonas*, and all metazoans (Fig. 2, Supplementary table S1). In alveolates and chlorophycean algae, the N-terminal region including h_1_ is divergent or absent, as documented by cryoEM structures of ATP synthases from *Tetrahymena* (Flygaard, et al. 2020), *Polytomella* (Murphy, et al. 2019) and *Toxoplasma* (Muhleip, et al. 2021). In *Polytomella*, subunit-a lacks the entire N-terminal region including the transmembrane helix h_1_, which is positionally replaced by the small transmembrane subunit Asa10 (Murphy, et al. 2019). However, Asa10 is unlikely to be a fragment of subunit-a, because it crosses the membrane in the opposite direction (Sinha and Wideman 2023). The missing helix h_N_ in *Polytomella* is positionally replaced by another conserved component, subunit-8 (Murphy, et al. 2019). Subunit-a in *Toxoplasma* is even more reduced and contains horizontal helices h_5_ and h_6_ only (Fig. 1C). On the contrary, subunit-a in *Tetrahymena* features a highly extended and structurally divergent N-terminus, which interacts predominantly with ciliate-specific ATP synthase components. Overall, the regions occupied by N-terminal parts of subunit-a in other ATP synthases are structurally highly divergent in *Toxoplasma* and *Tetrahymena*. The only alveolate with a canonical N-terminus of subunit-a is *Colponema* (Fig. 2), which supports the view that it represents a lineage sister to all other Alveolata (Tikhonenkov, et al. 2020).

In Metazoa, the N-terminal sequence of subunit-a differs from h_N_ in other groups, but invariably contains the motif FxxF (Fig. 2 and Supplementary table S1 and Supplementary Data 1). In experimentally determined and predicted structures of mammalian (Pinke, et al. 2020; Spikes, et al. 2020) and *Drosophila* subunit-a, respectively, this motif adopts the canonical fold of h_N_ (Fig. 1C), indicating that the ancestral architecture is retained despite sequence divergence. The only two exceptions are ctenophores and nematodes, represented by *Mnemiopsis leidyi* and *Caenorhabditis elegans*, respectively, whose subunit-a begins with helix h_1_, like subunit-a in trypanosomes. In *C. elegans*, this observation is in line with a recent cryoelectron tomography study that showed that the nematode ATP synthase diverged architecturally from canonical ATP synthases from yeasts and mammals and might lack subunits 8, k and i/j (Buzzard, et al. 2023), which are otherwise widely conserved in eukaryotes (Sinha and Wideman 2023). Due to high conservancy of h_N_ across eukaryotes, we wondered whether an equivalent of h_N_ could be retained in nematodes, possibly as a product of subunit-a gene split analogous to that observed in *T. brucei*. Such evolutionary independent event of fission of a mitochondrial gene would not be unprecedented. The gene encoding the mitoribosomal subunit uS3m has been fragmented in Ciliata (Swart, et al. 2012; Tobiasson and Amunts 2020) and some Heterolobosea (Fu, et al. 2014), and the gene encoding Cox2, a subunit of cytochrome oxidase, has been independently split at least three times in evolution, specifically in alveolates, chlorophycean algae (Perez-Martinez, et al. 2001; Waller and Keeling 2006; Rodriguez-Salinas, et al. 2012), and the hymenopteran insect *Campsomeris* (Szafranski 2017a). Therefore, we searched the proteomes of Nematoda using several approaches (blast, motif search, structure similarity search in AlphaFold database) to identify the protein constituting the missing region including h_N_, but we found no candidate, presumably due to the short length of the region and limited sequence conservancy.

In summary, the vast majority of eukaryotes with the exception of most Alveolata, a group of chlorophycean algae, and possibly Nematoda and Ctenophora have retained the ancestral architecture of the region corresponding to the N-terminus of subunit-a.

### Loss of transmembrane helix h_1_ predetermines the transfer of subunit-a from the mitochondrial to the nuclear genome

While the gene encoding subunit-a_N_ in Euglenozoa was transferred to the nuclear genome, the subunit-a gene has been retained in the mitochondrial genome. Likewise, subunit-a in nematodes, also starting with the first transmembrane helix h_1_, is encoded mitochondrially. In contrast, the gene of subunit-a in *Polytomella* and *Chlamydomonas*, which lacks h_1_, has been transferred to the nuclear genome. This suggests that while loss of h_1_ allows the transfer of the gene from the mitochondrial to the nuclear genome, the removal of the N-terminal fragment upstream of h_1_ is not sufficient for the relocation. Consistently, the reduced subunit-a in *Toxoplasma* and related species are also nuclear encoded (Huet, et al. 2018; Muhleip, et al. 2021; Fig. 2). The last known organism with subunit-a encoded in the nuclear genome is the ctenophore *M. leidyi*. The transmembrane helix prediction tools DeepTMHMM (Hallgren et al., bioRxiv), MEMSAT (Jones, et al. 1994), and HMMTOP (Tusnady and Simon 2001) do not predict h1 in the *M. leidyi* subunit-a. Alphafold does model h_1_ in *M. leidyi*, but with low confidence scores. Thus, it is unclear whether canonical h_1_ is present in the *M. leidyi* subunit-a or not. Together, these observations indicate that barriers associated with the insertion of h_1_-containing subunit-a to the inner mitochondrial membrane from the intermembrane space (or with translocation of the protein into mitochondria prior its insertion from inside) may be a factor limiting the transfer of the subunit-a gene from mitochondrial DNA to the nuclear genome. We expanded a previous analysis of hydropathy of subunit-a (Muhleip, et al. 2021) to all species included in our dataset. It revealed that some subunits-a (e.g., from *Nicotiana* or *Tetrahymena*) retained in mitochondrial genomes show relatively low hydrophobicity similar to those transferred to the nuclear genome (Fig. 2, Supplementary table S1), suggesting that the reduced hydrophobicity is not the only prerequisite for the intracellular subunit-a gene transfer. Regardless of the factor that allowed the subunit-a gene to be transferred, this is a very rare event that has most likely occurred only three times in the evolution of eukaryotes described to date, namely in the ancestors of the present-day myzozoans, ctenophores and Chlamydomonadales, an order of chlorophycean algae including *Polytomella* and *Chlamydomonas* (Fig. 2).

### h_N_ of subunit-a contributes to the coordination of the chain of proton translocating water molecules

The wide conservancy of h_N_ and its flanking residues (Fig. 2), whether within subunit-a or as part of a fission product, implies the functional importance of this region. The h_N_ contributes to coordination of the water chain in the lumenal proton half-channel in structurally characterized ATP synthases from *Trypanosoma* (Gahura, et al. 2022), *S. cerevisiae* (Guo, et al. 2017) and mammals (Pinke, et al. 2020; Spikes, et al. 2020) (Fig. 1B). One of the water molecules in the Grotthuss water chain is coordinated by positionally equivalent His19 of h_N_ of *T. brucei* subunit-a_N_ and Ser6 of h_N_ bovine subunit-a (Pinke, et al. 2020; Spikes, et al. 2020; Gahura, et al. 2022). In the vast majority of mitochondrial subunits-a, this position is occupied by another hydrogen bonding residue, glutamine, (Gln15 in *S. cerevisiae*; Figs. 1B&2 and Supplementary table S1). Thus, h_N_ most likely contributes to coordination of water molecules in most eukaryotic lineages and Alphaproteobacteria.

The shape of the inner access half-channel in bacteria resembles that of mitochondrial ATP synthase (Montgomery, et al. 2021), and a density consistent with presence of a water molecule in the inlet half-channel was observed in the cryoEM map of the *E. coli* ATP synthase (Sobti, et al. 2020). Early mutagenesis studies revealed several subunit-a residues crucial for enzymatic function of ATP synthase in *E. coli* (Cain and Simoni 1988; Lightowlers, et al. 1988; Vik, et al. 1988; Cain and Simoni 1989; Howitt, et al. 1990; Eya, et al. 1991; Hartzog and Cain 1994; Hatch, et al. 1995). Most of these residues are localized in the proton half-channels or at the interface between the subunit-a and the c-ring, implying their role in proton translocation. To assess to what extent the inner constricted portion of the lumenal proton half-channel differs between bacterial and diverse eukaryotic ATP synthases, we examined the conservancy of these residues and compared their positions with the residues proposed to be involved directly in the proton transfer or shown to coordinate ordered proton-transporting water molecules in mitochondrial ATP synthases (Spikes, et al. 2020; Gahura, et al. 2022); Fig. 1B&D, Supplementary figure 1). The only two invariable residues of subunit-a are the arginine of h5 separating the two half-channels and a glutamine of h6 (Arg210 and Gln245 in *E. coli* numbering), juxtaposing with each other. Asn214 and Glu196 lining the proton channel in *E. coli* are widely, yet not universally, conserved in mitochondrial ATP synthases. Functionally important His245 in *E. coli* is replaced by an acidic residue, in most cases glutamate (Glu203, Glu233 and Asp202 in bovine, *S. cerevisiae* and *T. brucei*, respectively), in vast majority of mitochondrial subunits-a. This residue is assumed to pass protons from the chain of water molecules to the c-ring either directly or by binding other, as yet unidentified, water molecules (Spikes, et al. 2020). Other functional residue, Glu219 in *E. coli*, is in most mitochondrial ATP synthases replaced by histidine (His168/155 in bovine/*T. brucei*), which binds one of the molecules in the water chain in the mammalian and trypanosomal enzymes (Spikes, et al. 2020; Gahura, et al. 2022). Thus, the positions of these two key histidine and glutamate residues are swapped between *E. coli* and mitochondrial ATP synthases. Nevertheless, due to the properties of these residues, both configurations are compatible with the Grotthus mechanism of proton translocation. Thus, the segment of the half-channel proximal to the c-ring most likely does not differ between bacterial and mitochondrial ATP synthases, supporting the view that the Grotthuss mechanism represents an ancestral and conserved mode of proton transfer in ATP synthases. In vast majority of mitochondrial ATP synthases and their alphaproteobacterial ancestors, h_N_ is a key component of the network of interactions involved in proton translocation.

### The entry point of the proton access half-channel in mitochondrial and alphaproteobacterial ATP synthases is defined by h_N_ of subunit-a

Notably, the *E. coli* ATP synthase (Sobti, et al. 2019) and ATP synthases in bacteria from various groups outside Alphaproteobacteria (*Mycobacterium;* Guo, et al. 2021; Montgomery, et al. 2021, and *Acinetobacter*; Demmer, et al. 2022) contain an α-helix at the N-terminus of the subunit-a, which is longer and sequentially dissimilar to h_N_ in mitochondrial ATP synthases and positionally shifted further from the loop connecting helices h_3_ and h_4_ (Fig. 1C). As proposed based on the structure of the *Acinetobacter* ATP synthase, the position of the N-terminal helix determines whether protons enter to the half-channel using the “front” or “rear” entry site as viewed from the c-ring (Demmer, et al. 2022). In *Acinetobacter*, the protons have been proposed to enter from the side facing the c-ring through the front entry located between the N-terminal helix and h_5_ of subunit-a. In *E. coli*, the protons are funneled from the periplasm to an entry point located between the N-terminal helix of subunit-a and the N-terminus of subunit-b (Yanagisawa and Frasch 2021), termed the rear entry (Demmer, et al. 2022). The aqueous funnel is lined with polar and charged residues, which are contributed by the N-terminal helix (Asp10, His14, His15) or h_1_ (Glu131, His132) and are not conserved in mitochondrial ATP synthases, and narrows to Glu219, presumably the first residue involved in the proton hopping. Notably, molecular dynamics simulations strongly suggest that protons can be delivered to Glu219 in *E. coli* also using another aqueous cavity (Ivontsin, et al. 2022), corresponding to the front entry in *A. baumanii*. Because *A. baumanii* and *E. coli* are closely related species and their subunits-a and adjacent regions are highly similar in both sequence and structure (Fig. 1, Supplementary data 1), we argue that both proton entry sites are most likely present also in *A. baumanii* (Fig. 3A).

**Figure 3.**
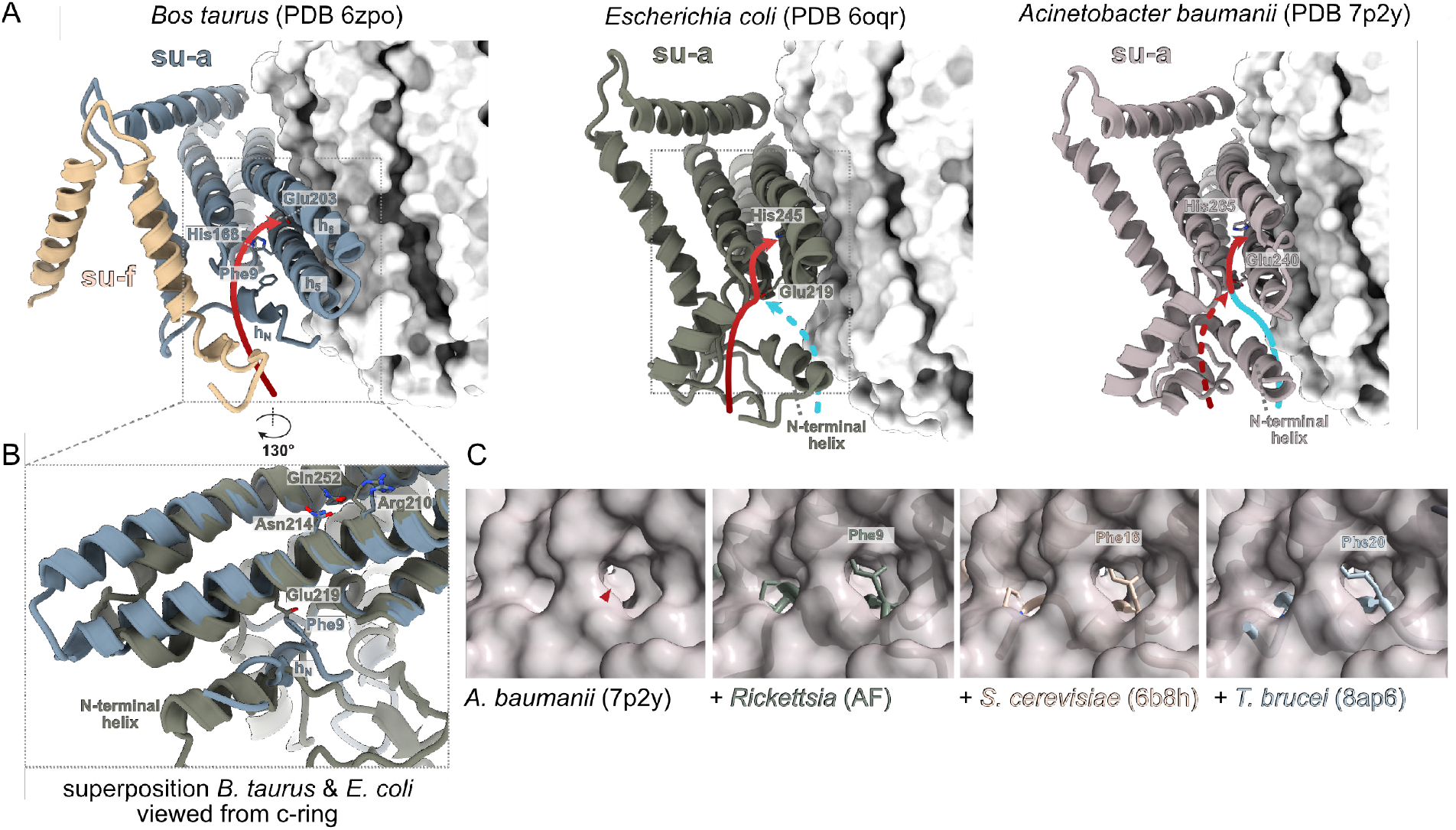
The proton entry point and path in the access half-channel is co-determined by h_**N**_. **(A)** Comparison of proton paths in the access half-channel of the bovine mitochondrial ATP synthases with *Escherichia coli* and *Acinetobacter baumanii* counterparts. The c-ring is shown as white surface. Blue and red arrows indicate the path through the front and rear entry, respectively. **(B)** Superposition of the structures from *E. coli* and bovine mitochondria shows that h_N_ is positionally shifted compared to the N-terminal helix of subunit-a in *E. coli* and that the conserved phenylalanine occupies the position of Glu219 in *E. coli*. Selected functionally important residues are shown as sticks in panels (A) and (B). **(C)** The position of the proposed front proton entry to the access half-channel in *A. baumanii* ATP synthase(Demmer, et al. 2022) is blocked by conserved phenylalanine in h_N_ of mitochondrial and alphaproteobacterial ATP synthases. A view of the front entry (red arrowhead) to the half channel in the *A. baumanii* subunit-a (surface representation) from the c-ring shown individually and superposed with subunit-a from other ATP synthases (cartoon representation): *S. cerevisiae*, the AlphaFold model of *Rickettsia*, and subunits a and a_N_ from *T. brucei*. The conserved phenylalanine is shown as sticks.

Unlike in *E. coli* and *A. baumanii*, in yeast and bovine mitochondrial ATP synthases protons use exclusively the rear opening, formed by a narrow cleft between h_N_ and the C-terminal tail of subunit-f(Spikes, et al. 2020; Demmer, et al. 2022) (Fig. 3A,B). In the yeast and bovine structures, as well as in the structure from *T. brucei* and the predicted structure from *Rickettsia*, the position of the front entry is blocked by h_N_ (Fig 3B,C), specifically by the phenylalanine residue of the SPLEQF motif that is nearly invariably conserved in all h_N_ sequences included in our analysis (Fig 2 and Supplementary table S1). Therefore, we reason that the rear entry is common to all h_N_-containing ATP synthases. In addition, the conserved phenylalanine clashes with the side chain of Glu219 in *E. coli* (Fig. 3B), which is the residue, at which the front and rear paths converge. Localization of coordinated water molecules in the part of the trypanosomal inlet channel that is more distal from the c-ring than Glu219 in *E. coli* (Fig. 1B) suggests that mitochondrial ATP synthases might use an extended water chain compared to bacteria.

Taken together, h_N_ contributes to shaping of both the surface-exposed and the inner portions of the access proton half-channel, underlying its almost universal occurrence in mitochondrial ATP synthases.

## Concluding remarks

By combination of analyses of available ATP synthase structures with sequence comparison and structure prediction across species, we documented that the proton access path and proton transfer mechanism in the access half-channel is conserved in majority of mitochondrial ATP synthases. The key structural determinant, so far largely overlooked, is h_N_ of subunit-a, which defines the entry to the access half channel and contributes to hydrogen bonding of water molecules in the Grotthuss chain (Fig. 1C). Unlike ATP synthase from *E. coli* and *A. baumanii*, which feature two possible proton entry sites, mitochondrial and alphaproteobacterial counterparts have only one access to the proton inlet half-channel, co-defined by h_N_ of subunit-a. Functional consequences of this difference remain to be elucidated.

Uniquely, F_1_F_o_-ATP synthase in *T. brucei* mitochondria contains two functionally central components, the subunits α and a, that are both constituted of two separate polypeptide chains. While the subunit-α is cleaved into two fragments posttranslationally (Gahura, et al. 2018; Montgomery, et al. 2018), the subunit-a is split on the gene level. Our finding of a split subunit-a in *T. brucei* complements recently reported cases of mitochondrial gene fragmentation revealed by structures of mitochondrial ribosomes (uS4m and uS7m in *Polytomella*; Tobiasson, et al. 2022) and respiratory chain supercomplexes (Cox3 and Nad5 in *Tetrahymena*; Mühleip, et al. 2022; Zhou, et al. 2022), and several occurrences reported earlier (reviewed in Szafranski 2017b). The transfer of the same gene from the mitochondrial to the nuclear genome independently in several lineages facilitates the inference of properties of encoded proteins that are incompatible with such events. We inferred that the transmembrane helix h_1_ might represent such an obstacle in the case of subunit-a.

## Material and Methods

Subunit-a sequences were searched by hmmsearch using the alignment of subunit-a sequences from *Bos taurus, Saccharomyces cerevisiae, Rickettsia rickettsii* and *Colponema vietnamica* as a query. The sequences from Ciliata and Apicomplexa were searched by phmmer using sequences from *Tetrahymena* and *Toxoplasma*, respectively, as queries. Both searches were performed on the HMMER web server (Potter, et al. 2018) in the Uniprot database. Amino acid sequence alignments were constructed by MUSCLE (Edgar 2004) and corrected manually using available structures as references. Some sequences of subunits a and a_N_ from Euglenozoa were obtained from (Sinha and Wideman 2023). Structures were predicted by AlphaFold (Jumper, et al. 2021), and superposed by the Matchmaker tool and visualized in ChimeraX (Goddard, et al. 2018). Hydrophobicity was calculated as grand average of hydropathy using hydropathy indices from Kyte and Doolittle 1982.

## Acknowledgements

We thank Anzhelika Butenko for the initial search of subunit-a homologs. This work was supported by Czech Science Foundation grant 20-04150Y to O.G. and by European Regional Development Fund (ERDF) and Ministry of Education, Youth and Sport (MEYS) project CZ.02.1.01/0.0/0.0/16_019/0000759 and by the European Research Council (ERC) under the European Union’s Horizon 2020 Research and Innovation Program (grant agreement no. 101044951) to A.Z.

## Author contributions

Conceptualization: O.G. and A.Z.; investigation and data analyses: J.E.W. and O.G.; writing – original draft: O.G.; writing – review and editing: J.E.W, A.Z., O.G.; funding acquisition: O.G. and A.Z.

## Declaration of interests

They authors declare they have no competing interests.

## Notes

### Competing Interest Statement

The authors have declared no competing interest.

### Summary of Updates

The manuscript has been revised based on a peer-review in a journal.

