## Supplementary figure S1 for "The ancestral shape of the access proton path of mitochondrial ATP synthases revealed by a split subunit-a"

**List of supplementary material:**

**Supplementary table 1:** Features of subunits a and $a_N$

**Supplementary data 1:** Sequences of subunit-a from eukaryotes

**Supplementary data 2:** Sequences of subunit-a from alphaproteobacteria

**Supplementary data 3:** Sequences of subunit-$a_N$ from Euglenozoa

**Supplementary figure 1.**

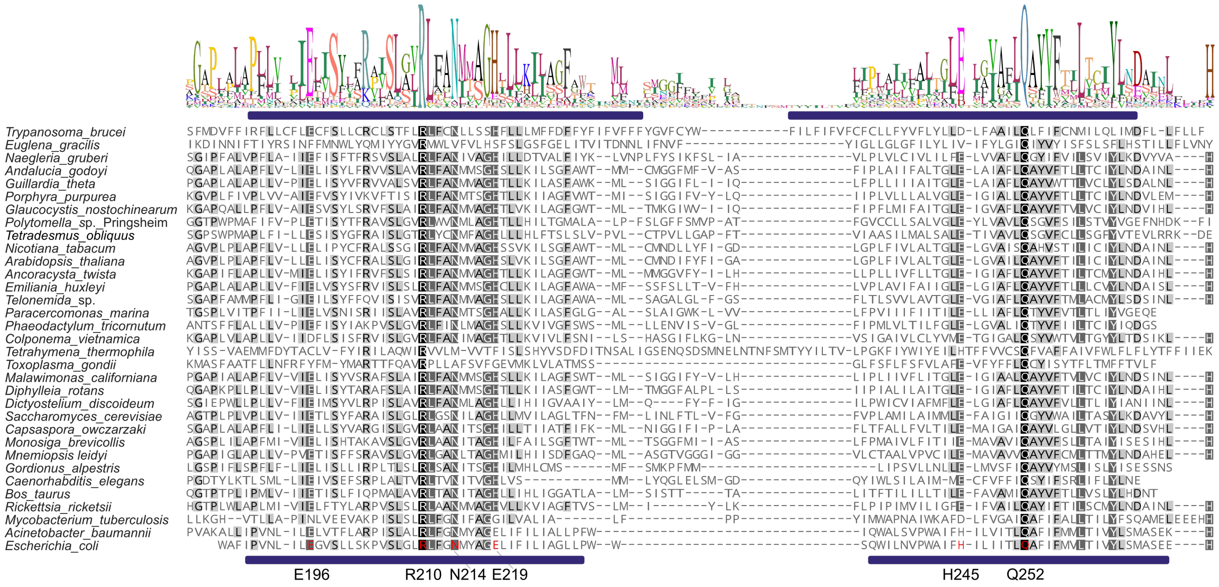


**Supplementary figure 1. Sequence alignment of the region of subunit-a encompassing helices h$_5$ and h$_6$ from selected organisms.** Blue bars indicated h$_5$ and h$_6$ sequences from *T. brucei* and E. coli. Selected *E. coli* residues are highlighted.
